# Genetic drift from the out-of-Africa bottleneck leads to biased estimation of genetic architecture and selection

**DOI:** 10.1101/2020.08.17.254110

**Authors:** Bilal Ashraf, Daniel John Lawson

## Abstract

Most complex traits evolved in the ancestors of all modern humans and have been under negative or balancing selection to maintain the distribution of phenotypes observed today. Yet all large studies mapping genomes to complex traits occur in populations that have experienced the Out-of-Africa bottleneck. Does this bottleneck affect the way we characterise complex traits? We demonstrate using the 1000 Genomes dataset and hypothetical complex traits that genetic drift can strongly affect the joint distribution of effect size and SNP frequency, and that the bias can be positive or negative depending on subtle details. Characterisations that rely on this distribution therefore conflate genetic drift and selection. We provide a model to identify the underlying selection parameter in the presence of drift, and demonstrate that a simple sensitivity analysis may be enough to validate existing characterisations. We conclude that biobanks characterising more worldwide diversity would benefit studies of complex traits.

## Introduction

Understanding complex traits is one of the most important questions facing genetics as we progress into the Biobank era. The number of Single Nucleotide Polymorphisms (SNPs) that influence complex traits may vary from tens to thousands in human and non-human species (1,2). The effect of each SNP on a trait is estimated using Genome Wide Association Studies (GWAS) in the very large biobanks and meta-analyses needed for statistical power. Because of the requirement for large sample sizes, almost everything that we know comes from studies in Eurasia in which these datasets are available; for example the UK Biobank (3), the China Kadoori Biobank (4), the Japanese Biobank (5) and large GWAS consortia (6,7). Yet, most selection acting on complex traits occurred primarily in our evolutionary history. How did the out-of-Africa bottleneck (8) influence our quantification of complex traits?

There is much interest in describing the genetic architecture (9) of complex traits. If a trait is under negative or balancing selection, then SNPs with a large effect are selected against, and reduced in frequency. Genomic (or Genetic) architecture quantifies the relationship between SNP frequency and the effect the SNP has on the trait (10). Models (11,12) use an explicit parameter that we will denote *S* that describes this shape, and which is often linked to selection. *S* = 0 means that effect size and SNP frequency are unrelated. *S* < 0 means that rare SNPs have larger effect, and is expected if large effect SNPs are driven to low frequency by negative or balancing selection. Conversely, *S* > 0 implies that common SNPs have a larger effect, and is expected if selection increases the frequency of large effect SNPs via positive selection.

Genetic drift (13) is the process of SNPs varying in frequency over time due to individuals carrying the SNP having a random number of offspring each generation. It is well understood in a nearly-neutral context (14) allowing for limited selection. Clearly, the genetic architecture representation as a conditional model describing the effect size, conditional on the SNP frequency, is incomplete. Whilst the allele frequency spectrum is related to selection (15), a joint model is much more difficult, especially when ascertainment, linkage and other statistical artefacts are accounted for. Figure 1 illustrates how Genetic Drift and Complex Trait Genetic architecture interact to change the whole SNP-frequency and effect size distribution.

**Figure 1:**
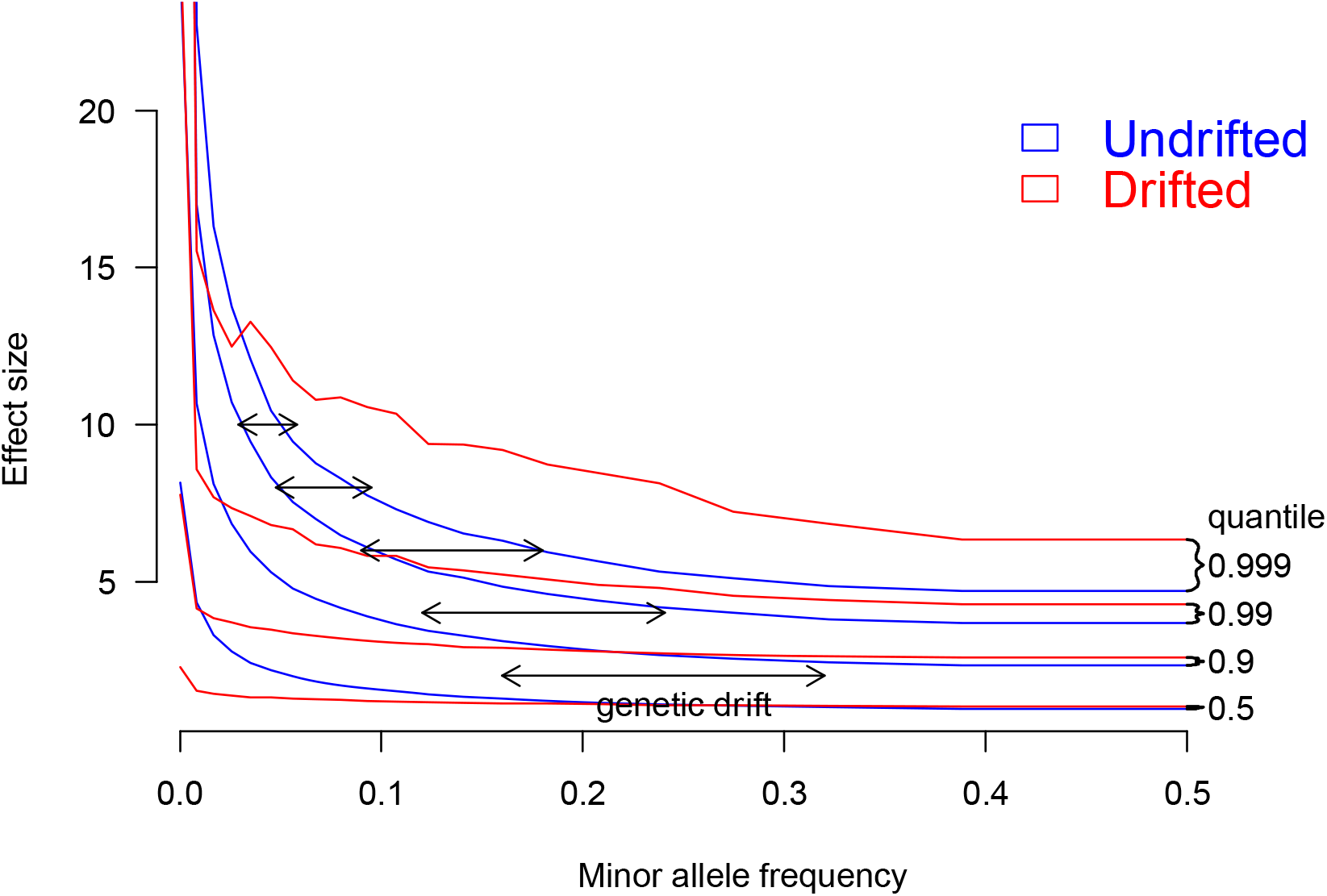
Simulation of Complex Trait Genetic architecture with Genetic Drift. The Complex Trait has *S* =−1, meaning that most large effect alleles are very rare. The blue distribution shows quantiles of effect size in the population in which the trait evolved, conditional on frequency. Genetic drift (here, *F_st_* = 0.1) changes the blue to the red distribution. Drift is larger for common SNPs with modest effect, so most rare SNPs either become a little more common, or go to fixation. The result is a much flatter distribution (e.g. the 0.5, 0.9, 0.99 quantiles) which resembles a smaller magnitude shape parameter *S*. However, the most extreme SNPs at a given frequency (q=0.999) arrive from lower frequency and hence have much larger effect. Whilst the red distribution cannot be exactly replicated by a different shape parameter *S*, it can be closely approximated if relatively few SNPs contribute to the complex trait.

We use a simulation approach to examine whether the out-of-Africa bottleneck should change the interpretation of parameters in the genetic architecture of complex traits. We find that inference in a *target population* of Europeans, and any other non-African population have a rather different genetic architecture to the *evolving population*, proxied by Africans, in which selection predominantly occurred. As a consequence, *S* cannot be understood as a direct quantification of selection, and indeed the value obtained depends on many things including any Minor Allele Frequency (MAF) thresholding performed in quality control. Models of genetic architecture that do not correct for drift are a useful description of the data, but further work is needed for inference about selection.

## Results

### Genetic architecture is changed by genetic drift

#### Simulation framework

To assess the effect of genetic drift on genetic architecture we need a large sample of individuals from around the world, which is not currently available. To address this we resample data from the 1000 Genomes dataset (16) using HAPGEN2 (17) to create realistic population structure complete with linkage disequilibrium between Africa, Europe, South Asia, East Asia, and America. We then simulate complex trait effect sizes in the African population (see Methods). To generate individual data, we use (narrow sense) heritability *h*^2^ = 0.5 throughout. We vary the SNP frequency relationship *S*; recall that *S* < 0 implies “negative selection” on the trait, and therefore high frequency SNPs can only have a small effect on the trait, whilst rare SNPs are permitted to have larger effect sizes.

To generate genetic variability in each of our populations we follow (18) by assuming a relationship between frequency *f* and effect size *β*, for each SNP *i* of the form:

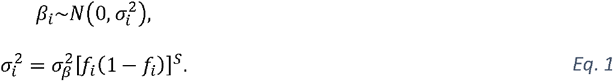

where 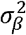 is a base-rate variance (see Methods). However, the details depend crucially on how variants that are rare in the *evolving population* are treated in the generative model for complex traits. Because less information is available about real rare variation, little is known about how, in reality, these affect complex traits. One reasonable assumption is that the effect size follows the model described above for all frequencies *f_i_* (the default *unbounded effect simulation*). However, this leads to rare SNPs having unbounded effect size. An alternative reasonable assumption is that 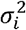 is bounded (referred to as the *bounded effect simulation*) (Methods).

Heritability is the proportion of variance attributable to genetic variation, and therefore depends critically on assumptions about transferability of environmental variation. Our simulation assumes a constant value for environmental variability, determined to be that required in Africans to give *h*^2^ = 0.5 with the specified MAF threshold, from which we compute an observed heritability *h*^2^. We also report values computed with GCTB using --bayes S (12).

Finally, in real data analysis, it is necessary to exclude SNPs that are very rare in the *target population* by excluding those beneath some MAF threshold. These need not be the same SNPs that were rare in the *evolving population*.

#### Inference

The resulting heritability for simulated complex traits in African and other populations is given in Figure 2. Both our approach and GCTB agree that heritability in non-Africans is strongly biased by the bottleneck, and that the magnitude of this effect is a function of the simulated value of *S*. However, we observe that thresholding critically impacts the inferred heritability. If no thresholding is performed, the inferred *h*^2^ is significantly larger than simulated, whilst if thresholding is strict, the inferred *h*^2^ may be smaller. It is not that this heritability is “wrongly estimated”, but is a property of a trait realised in a specific population due to different genetic variation leading to different phenotypic variation.

**Figure 2:**
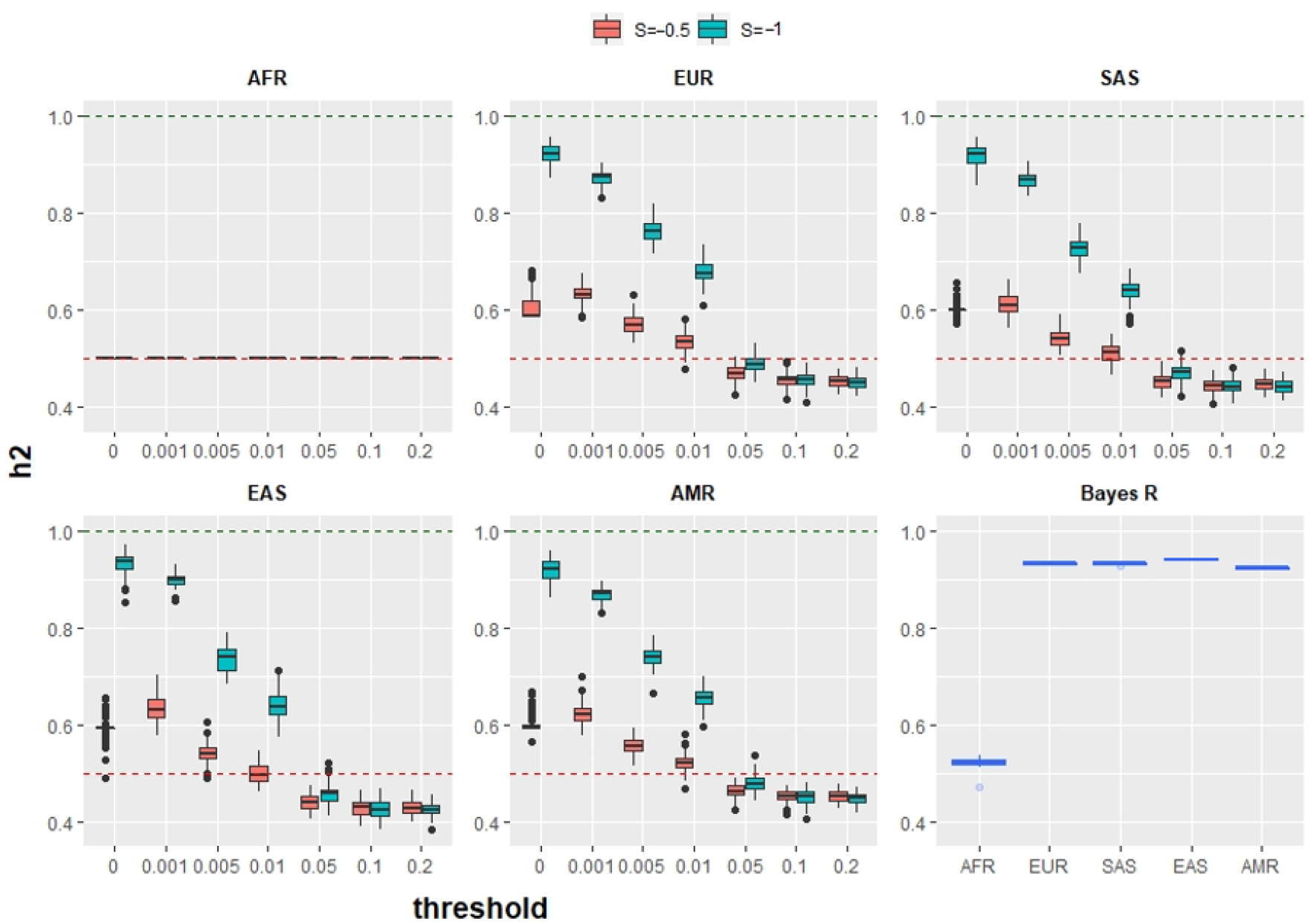
Estimates of heritability when a complex trait is simulated in 1000 Genomes Africans (AFR) with *h*^2^ = 0.5 and observed in any other worldwide population, when environmental variability constant across all populations. Each plot shows observed heritability at different thresholds for SNP frequency, for a different population group at *S* = −0.5 and *S* = −1. The final plot (Bayes *S* panel) shows results from GCTB --bayes R (30) for *S* =−1, which agree with our unthresholded estimates.

This is a direct consequence of genetic drift changing allele frequencies independently of SNP effect size (Figure 2). Low frequency SNPs with large effects can become common, leading to an increased genetic variation of the trait (Supplementary Figures 1–2). This is precisely why bottlenecked populations including Ashkenazi Jews (19), Finns (20) and Icelanders (21) are used in GWAS studies for generally rare diseases that are common in those populations.

It is important to emphasise that these heritability changes are a consequence of the total genetic variation changing as a consequence of genetic drift. Similarly, environmental variation for real phenotypes varies due to factors including lifestyle, societal organisation, and so on. We report these heritability results to emphasise how important assumptions are in modelling. Of course, it is possible to scale the environmental variation with the genetic variation to ensure a desired heritability. Changing the environmental variation added to phenotypes will not affect the inference that follows.

### Inferred selection is affected by genetic drift

We then asked whether the relationship between SNP frequency and effect size has been distorted by genetic drift, by estimating the selection coefficient *S*. For this we implemented Eq. 2 as a Bayesian model (see Materials and Methods).

We call this the “simple model” as it does not account for genetic drift. This relationship is typically a prior that affects effect size estimates; for our model this is a likelihood for the observed effect size, which we assume given. These would be taken from GWAS, but in simulations effect sizes are treated as known. This eliminates the estimation error that often dominates genetic architecture studies.

Figure 3 shows that, like, is biased by genetic drift, but this depends critically on how the phenotype is truly formed. In the *unbounded effect simulation* where no MAF thresholding is performed (threshold=0.0001 excludes only SNPs absent in Africa), the inferred is larger in magnitude than the simulated. Conversely, in the *bounded effect simulation* can be below the true value, and for large thresholds tends towards the prior mean of 0, due to a lack of variation in the data. There is a transition around minor-allele-frequency of 0.05 where the biases cancel out. However, there is significant variability in the inferred, due to the random nature of genetic drift and the sensitivity of the inference to the most extreme causal SNPs.

**Figure 3:**
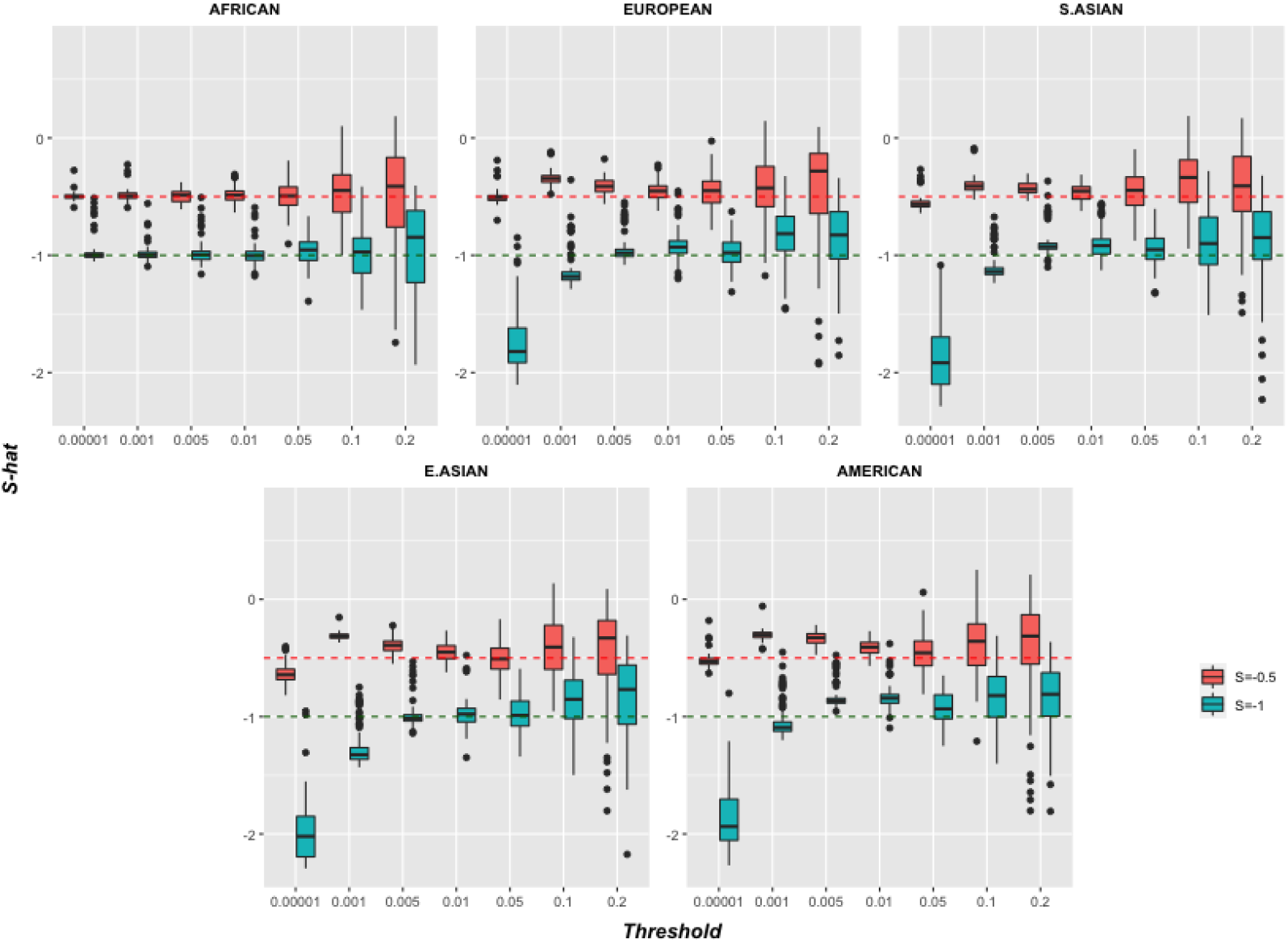
Inferred architecture parameter *S* with different thresholds for all 1000 Genomes population groups, using a simulated = −0.5 and = −1. The complex trait was simulated in Africans and inferred in the specified population using the “Simple model”. See Methods for details.

Unlike for heritability, it is not clear how a simulation should be updated to maintain a desired *S*. The choice of environmental variation does not effect *S* as it is simply adding different amounts of noise to the phenotype. This is therefore a rather different sort of bias.

Critically however, the choice of MAF thresholding does not affect inference in the population that experienced the selection; in our simulations this is Africa (AFR). In this population, accurate estimates of *S* are recovered for a range of thresholds (up to MAF 0.1, above which power is lost) which induced considerable bias in every other population. MAF thresholding is therefore a potential sensitivity analysis tool for the interpretation of *S*.

### Separating drift and selection

Bias in heritability and *S* are both natural consequences of genetic drift. To model genetic drift and hence recover the pre-drift values (see Materials and Methods) we allow for genetic drift in a “drift model” (Figure 4), in which the drift process is represented using the Balding-Nichols model (22). As no individual data is required, these simulations are larger (N=4000) than Figures 2–3 (N=1000). We demonstrate two cases where the drift model works well; when the Balding-Nichols model holds (Figure 4a,d) and also in the *bounded effect simulation* (Figure 4b,e,f) where it may approximately hold. In these cases, significant under-estimation of *S* is observed in the simple model that ignores drift, which grows with true *F_st_* (Figure 4d). We also demonstrate the requirement to accurately estimate *F_st_* (Figure 4f) in the appropriate SNP set; use of genome-wide non-representative estimates can create bias in the drift model.

**Figure 4:**
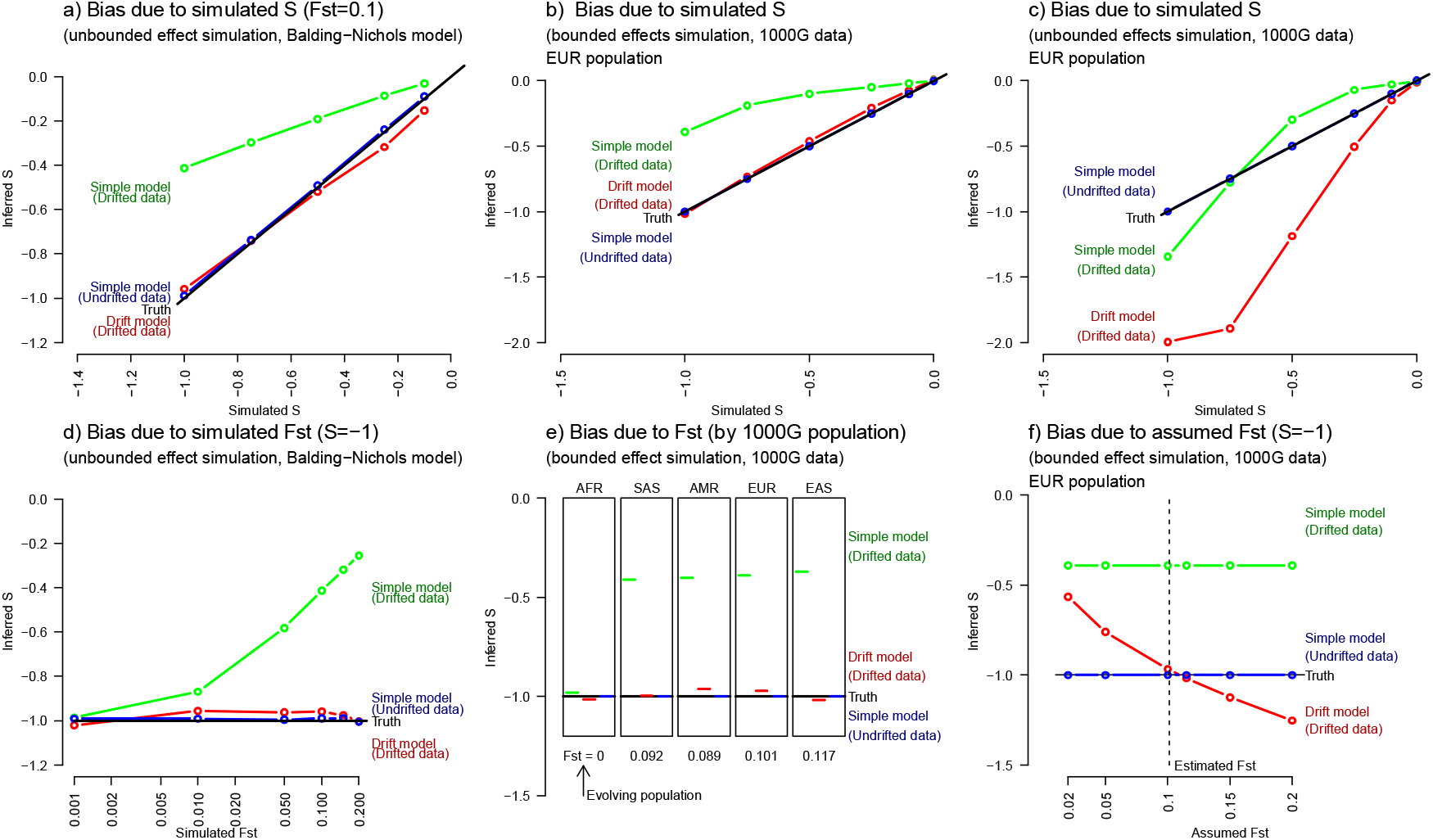
Drift aware inference of genetic architecture removes bias if the model holds. a) Simulation of genetically drifted genetic data in the unbounded effect simulation using the Balding-Nichols model of genetic drift as used for inference leads to biased estimation of genetic drift in the “simple model” which is corrected by our drift model. b) The same bias is observed for simulation of genetically drifted genetic data in the bounded effects model in the European (EUR) population, which is corrected by our “Drift model” (estimated *F_st_* = 0.101 in these SNPs, see Methods). c) However, where SNPs with large effect that do not fit the drift model are included (the unbounded effects model) all inference is biased. d) In the Balding-Nichols simulation we can vary genetic distance *F_st_* and find a nonlinear relation. e) The bias can be corrected for all 1000 Genomes populations with phenotypes generated in Africa (AFR), and examined in South Asians (SAS),Native Americans (AMR), Europeans (EUR) or East Asians (EAS). f) The estimate is sensitive to the estimate of *F_st_*, which must be performed on data representative of the included SNPs. (Plots show median and 90% credible sets for inference and all use “Africans” to proxy the evolving population.)

Unfortunately, due to model mis-specification, the drift model fails in the *unbounded effect simulation* of 1000 Genomes data in which the rare SNPs with high effect size are not correctly modelled (Figure 4c), leading to bias in all considered models, for any population except the evolving one.

The bias in *S* is controlled by two competing effects. *S* is inferred to be larger in magnitude if genetic drift takes rare SNPs with large effect to high frequency, where they are unlikely according to the model without drift and therefore dominate the inference. The number of such SNPs increases rapidly with *S* (Supplementary Figure 3), requiring fewer than 10k SNPs with *S* = −1, 1M SNPs with *S* = −0.5, and more than we could simulate with *S* = −0.25. Conversely, without SNPs with large effect, genetic drift leads *S* to be closer to 0 as effect sizes become more homogenous.

Complex Trait architecture is affected by any MAF frequency change, not just genetic drift. Archaic admixture (see Methods) has led to a small fraction (around 5%) of SNPs with high differentiation (from Neanderthals for all non-Africans cite (23) and Denisovans (24) in Oceania and South East Asia. Simple simulations show (Supplementary Figure 4) that archaic admixture is not likely to be the largest contributor to high frequency SNPs, as genetic drift from the out-of-Africa event is already capable of creating dramatic changes in SNP frequency. However, this is assuming nearly neutral drift of individual SNPs, i.e. that genetic drift dominates selection at the individual SNP level. Loci selected out of modern humans due to large effects do not fit the “genetic architecture” framework and could be introduced via archaic populations.

## Discussion

Selection occurred on most complex traits in the evolution of modern humans; that is, most selection will have acted on the *evolving population* prior to the out-of-Africa event that led to the peopling of Eurasia and beyond. This bottleneck led to considerable genetic drift in all non-Africans, which can bias inference of selection where these are used as a *target population*.

What do our results imply for real complex traits? Unfortunately, little can yet be stated with confidence where traits have been analysed without consideration of drift. We demonstrated that the bias in *S* can be positive or negative, sensitively to details of complex traits that are not currently well understood: the true value of *S* and the effect sizes of SNPs that were rare in the evolving population. Inferred *S* is more extreme in drifted populations if the effect size of extremely rare SNPs is appropriately modelled by the bulk of the distribution. However, it is smaller if the effect size remains bounded. From an “extreme value” perspective (Supplementary Figure 4), we hypothesise that the presence of a small number of SNPs with strong effect coupled with much missing heritability is an indication of being in the “under-estimated *S*” regime.

We hypothesise similar issues surrounding model-misspecification of the complex trait. For example, if the distribution of effects is not normal, if the variance does not fit the assumed model, or if typical non-ancestral variants have a biased effect (e.g. are weakly maladaptive). In such a misspecified model, details such as the prior on the noise can affect inference; for example, the scale of the variation in effect size (*σ_β_*) may matter. It is likely that semi-parametric models, which are not sensitive to the distribution of effect sizes in the bulk of the SNPs, will be more robust to these issues, and potentially restricting inference to common SNPs in both Europeans and Africans will aid robustness.

More constructively, we demonstrated that a simple sensitivity analysis, that of performing inference at a range of minor-allele frequencies, can identify whether genetic drift has an influence on the inferences made on a particular complex trait. We then showed that correcting for genetic drift was plausible and desirable, and provided a Bayesian inference algorithm for this.

It is important to emphasise that our algorithm implements the “prior” component of the model and can only be used on real data if unbiased estimates of effect sizes (allowing uncertainty) can be obtained. Whilst our implementation lacks the SNP selection component of established tools, our model can be directly used by performing SNP selection within other software, or software could be updated to allow more appropriate models. *S* is always a valid summary of a specific genetic architecture, but to link *S* to selection it is essential that sensitivity analysis or further modelling supports this interpretation.

Our model uses relatively little information and is not likely to reconstruct true allele frequencies from the past; it instead learns ancestral SNP frequencies that make the Complex Trait effect size distribution most plausible. It also does not implement inference of *F_st_*, as it would be inconsistent to infer *F_st_* on a trait-by-trait basis for the same SNP set. However, it is the case that *F_st_* varies considerably between SNP sets and the *F_st_* we observed across populations was low, which may be due to the relatively high frequency imposed on this during SNP selection.

Genome-Wide, *F_st_* between Africans and Eurasians is high at ~0.2 (16); within Eurasians is moderate (~0.1 between Europe/China) and small within ancestry groups (~0.01 between North and South Europe). Yet the appropriate *F_st_* from the ancestor of all humans is not completely clear. Diversity within Africa is extremely high (again ~0.2-0.3) (25). As larger datasets within Africa become available, we will need to establish whether selection has continued to operate effectively on complex traits, leading to unbiased estimates from these populations. If not, it may still be inappropriate to use a specific modern African population as a proxy for the ancestral population of modern humans. Despite this, African individuals who have not experienced the bottleneck will be essential in establishing the true genetic architecture of complex traits, as drift modelling alone will have limited power to infer the original SNP frequencies.

On Complex Traits whose variation is dominated by relatively few SNPs, it will be hard to separate genetic drift and selection. This leads to two independent avenues of further research. The first is to increase diversity of large-scale population studies and especially African ancestry, to access the genetic diversity that was lost in the Out-of-Africa bottleneck. The second is to develop multi-ethnic models of genetic architecture to account for population structure.

## Materials and Methods

### Data sets

#### The 1000 Genomes Project

We use the 1000 Genomes Project data for simulation analyses. The latest release is phase 3, containing 84.4 million variants for 2504 individuals. Population groups in this data are African (AFR), European (EUR), South Asian (SAS), East Asian (EAS) and American (AMR) (16).

1000 Genomes data (genome wide) were pruned based on linkage disequilibrium. Variant pruning was done using PLINK 1.9 (26,27) with command LD--indep-pairwise 200 10 0.07. After pruning 354,443 SNPs were retained. These SNPs were further passed to HAPGEN2 (17) to simulate 10,000 individuals from each population. The data set for analysis was 10,000 individuals, 354,443 SNPs for each of five population groups were available for further analysis.

#### Complex trait simulation

We generate a random complex trait by selecting *N* causal SNPs at random, and simulating effects from our model following (18): 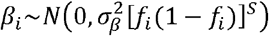. We set 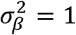 without loss of generality, and *f*_i_ are taken as the African SNP frequencies.

Individual level data are required for running GCTB, and for the computation of heritability and genetic variance under genetic drift. Then the genetic variation 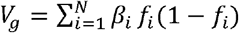, and we fix narrow sense heritability(28) 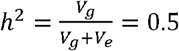 in the evolved population to set the environmental variation *V_e_* = *V_g_*(*evolved*). we use *N*=1000 and the phenotype of an individual *k* is sampled from their (binary) genome *x_ki_~Bern*(*f*_i_) as the sum of genetic plus environmental contributions 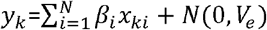. All 354,443 SNPs were passed to GCTB, but only the *N* causal SNPs were considered by our algorithm. For Figure 4 in which no individual data is generated, *N*=4000.

#### Bayesian model for Genetic architecture with drift

We created a novel MCMC algorithm in Stan (29) (mc-stan.org) using the Rstan interface.

Model 0 is the baseline model which is an implementation of the BayesS model in which there are no SNPs that do not affect the trait, because we know which these are. Model 0 can be written for each SNP *i* = 1.. *L* for the observed frequency *f*_i_ and observed effect size *β*_i_:

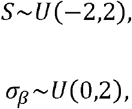

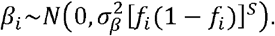

The “drift model” is an extension accounting for genetic drift. It follows Model 0, except that we simulate the complex trait in a “pre-drifted population”. SNP frequencies in this population is *p_i_* which generates the “drifted data” frequency *f*_i_ using the Baldings-Nichols model (22) to represent drift using the “Fixation Index” *F_st_*, treated as known. This leads to:

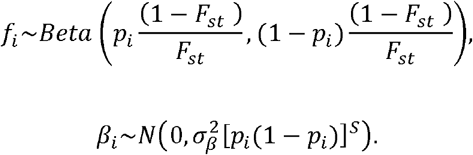

Here, Normal distributions are specified via (mean, variance) and the Beta distribution is specified as *Beta(α,β)* defined in terms of shape and scale parameters with expectation *α*/(*α + β*). Therefore *f*_i_ has expectation 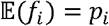, and variance Var(*f*_i_) = *F_st_ p_i_*(1 − *p_i_*).

When *F_st_* is known (Figure 4a-b) this is provided to the model. When *F_st_* is unknown, we estimate it on our dataset using plink1.9 (www.cog-genomics.org/plink/1.9/) (26) using “--fst –within”, providing only the individuals belonging to the two populations being compared.

Unless otherwise stated, all SNPs are considered without thresholding in the *target population*, except for those that have reached fixation which are omitted as they have zero probability under the likelihood.

For Figure 4 we run 10 replicates using 4 chains each and retain only runs that converged according to the *Rhat* statistic (29) using the criterion *Rhat(S)*<1.2. Typically, each chain either converges rapidly to the correct mode (*Rhat*<1.02 in 78% of replicates) or one or more chains become stuck in a poor local optima with S>0 leading to Rhat >=1.5.

#### Default and bounded effect simulation for effect sizes

The difference between these models is created solely by the selection of SNPs to be included in the simulation. For the *default simulation*, all SNPs with frequency >0 in Africans are considered. For the *bounded effect simulation*, only SNPs with frequency >0.01 in Africans are considered for sampling.

The nomenclature arises from the consequences of this thresholding. The variance of SNPs in the *bounded effect simulation* is therefore bounded at [*p_i_*(1 − *p_i_*)]*^S^* ≈ 101 if *S* = −1 and *p_i_* = 0.01; compared to a minimum variance of 4 if *p_i_* = 0.5. This is 200 times smaller than the variance of 20001 assigned to the rarest SNP in the dataset (*p_i_* = 5*e*^−5^).

#### Simulation model for Supplementary Figure 3

We created a simulation model that could characterise our model rapidly without going through the 1000 Genomes data, hence providing a simulation that could generate a range of simulated *F_st_* values and demonstrating performance under the assumed model. We choose a value of *S* and *F_st_* and then simulate data from the “drift model” with a specified *L* (=10000 throughout).

We also threshold MAF to 0.0001, i.e. in the inference model, any frequency less than 0.0001 is treated as 0.0001.

## Acknowledgements

DJL and BA were funded by the Wellcome Trust under grant number WT104125MA.

## Code availability

The code necessary to replicate the results presented here are given at https://github.com/danjlawson/genomicarchitecture.

## Supplementary Figures

**Supplementary Figure 1:**
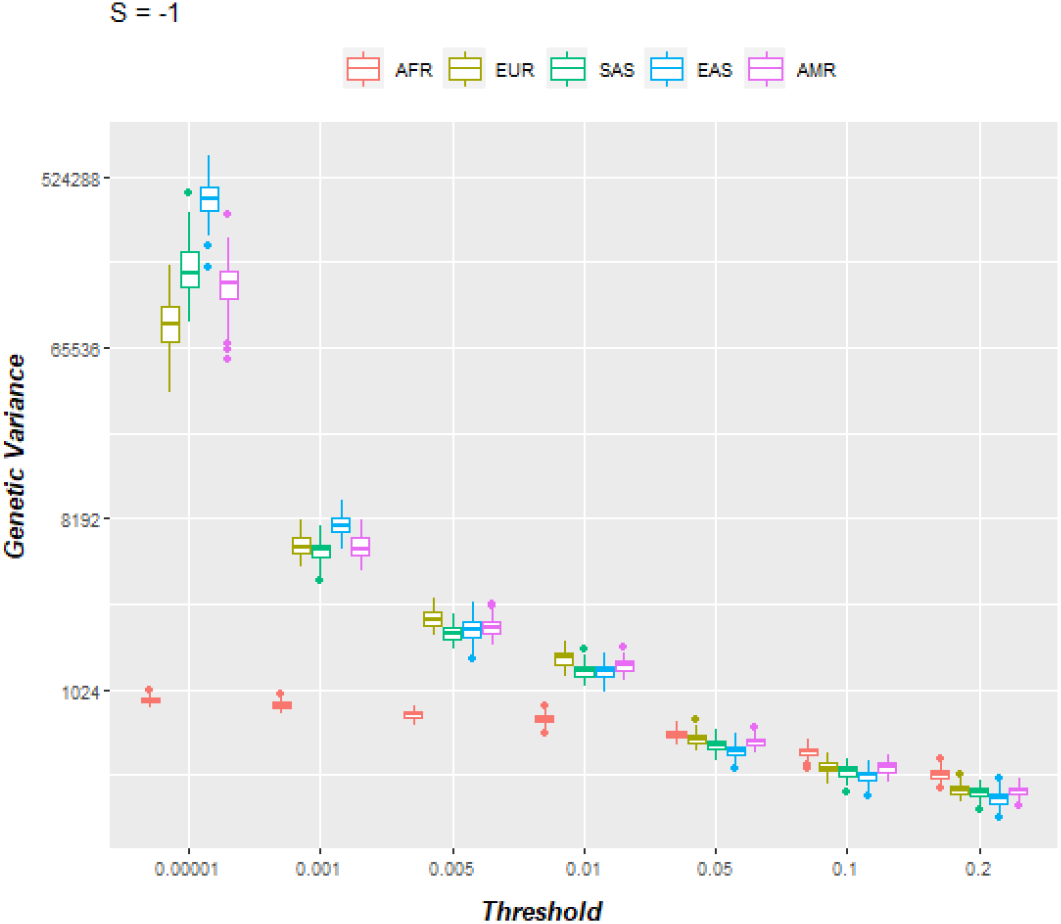
Estimates of genetic variance with different thresholds for all population groups at *S* = −1, as reported in Figure 2 in the form of Heritability.

**Supplementary Figure 2:**
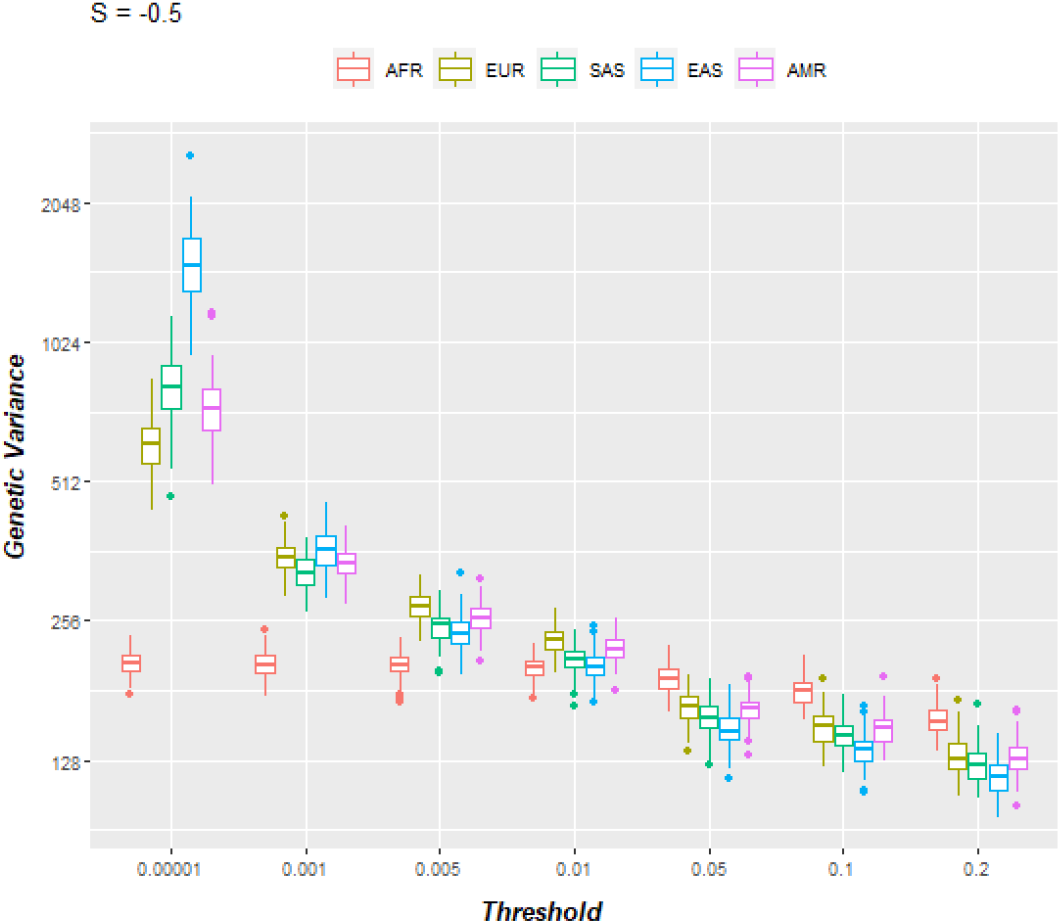
Estimates of genetic variance with different thresholds for all population groups at *S* = −0.5, as reported in Figure 2 in the form of Heritability.

**Supplementary Figure 3:**
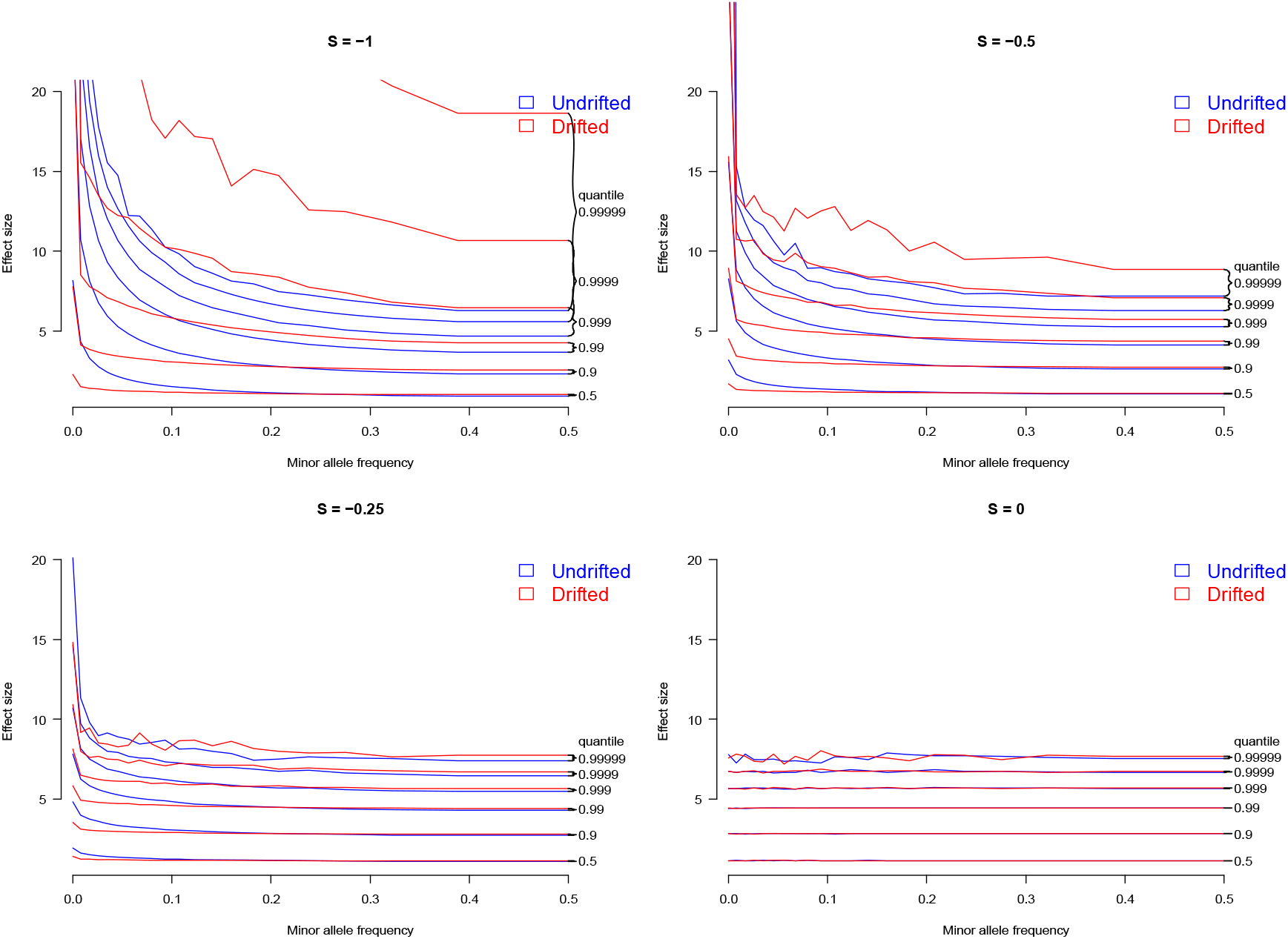
Simulation of Genetic architecture with Genetic Drift at varied values of *S* (Assuming *F_st_* = 0.2). The model and presentation follows Figure 1, and varies only by a) showing a larger range of quantiles, and b) shows *S* =−1,− 0.5,−0.25,0. The number of SNPs required for drift to result in a large effect SNP that dramatically changes model inference is of the order 1/(1-q) where q is the first quantile for which the drifted model has significant mass above the corresponding undrafted curve.

**Supplementary Figure 4:**
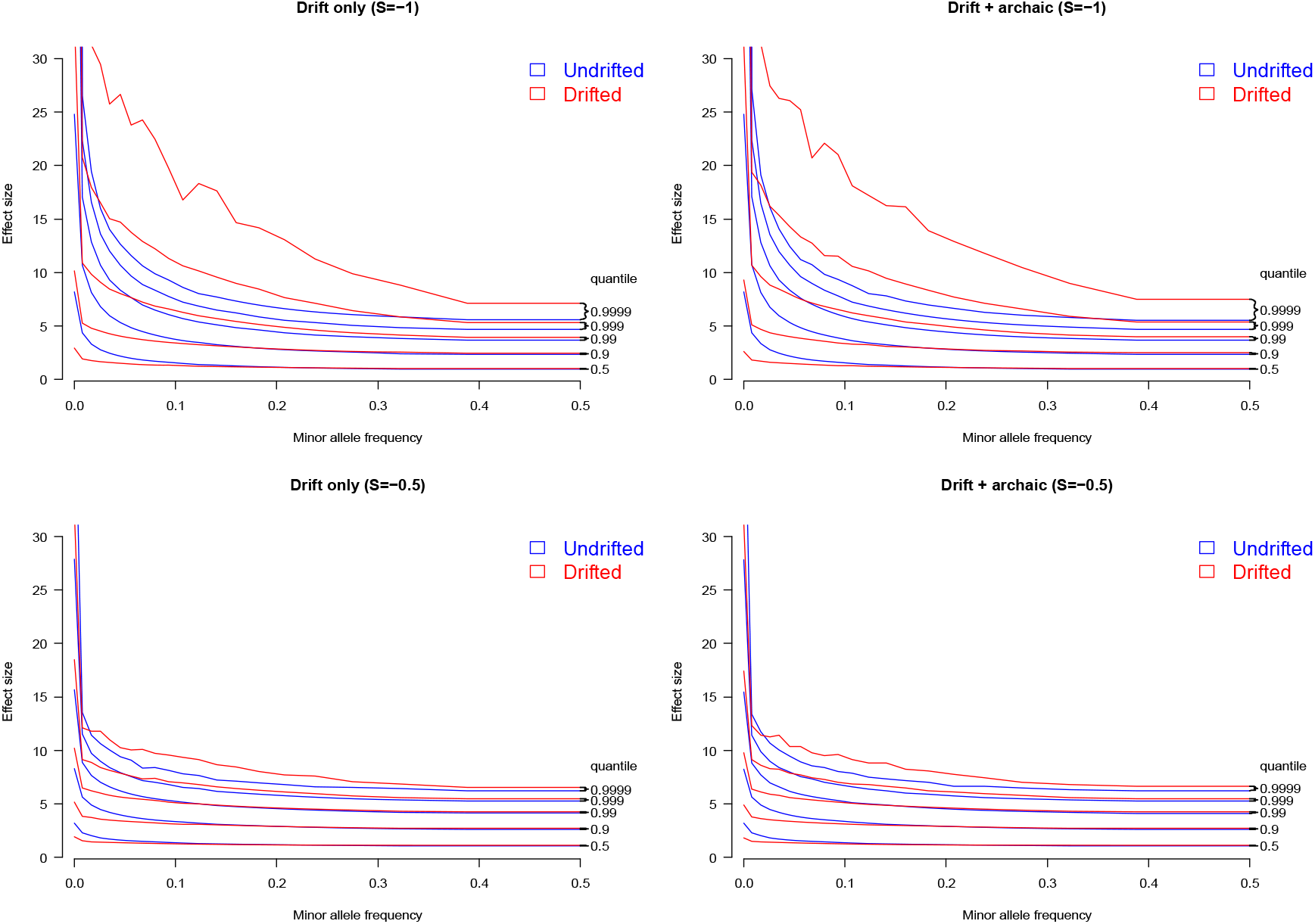
Impact of archaic admixture on Genetic architecture, assuming *F_st_* = 0.1, comparing (left) Genetic drift from the Out-of-Africa event only, to (right) additionally adding 5% of the genome having *F_st_*= 0.5.

## Notes

https://github.com/danjlawson/genomicarchitecture

### Competing Interest Statement

The authors have declared no competing interest.

